# Identification of PCNA interacting protein motifs in human DNA polymerase delta

**DOI:** 10.1101/2020.02.29.971473

**Authors:** Prashant Khandagale, Shweta Thakur, Narottam Acharya

## Abstract

DNA polymerase delta (Polδ) is a highly processive essential replicative DNA polymerase. In humans, Polδ holoenzyme consists of p125, p50, p68, and p12 subunits and recently, we have shown that p12 exists as a dimer. Extensive biochemical studies suggest that all the subunits of Polδ interact with the processivity factor proliferating cell nuclear antigen (PCNA) to carry out a pivotal role in genomic DNA replication. While PCNA interaction protein (PIP) motifs in p68, p50 and p12 have been mapped, the PIP in p125, the catalytic subunit of the holoenzyme, remains elusive. Therefore, in this study by using multiple approaches we have conclusively mapped a non-canonical PIP box from residues _999_VGGLLAFA_1008_ in p125, which binds to inter domain-connecting loop of PCNA with high affinity. Collectively, including previous studies, we conclude that similar to *S. cerevisiae* Polδ, each of the human Polδ subunits possess motif to interact with PCNA and significantly contribute towards the processive nature of this replicative DNA polymerase.

## 1. Introduction

Eukaryotic DNA replication requires the concerted action of several enzymes and accessory cofactors [1–3]. While helicases and DNA polymerases are the key enzyme components; replication factor C (RFC), proliferating cell nuclear antigen (PCNA) and replication protein A (RPA) are the integral structural components of the DNA replication machinery. Thus, these genes are essential for cell survival. Again, interestingly, all these factors are multi-subunits. Since, the two strands of DNA are in antiparallel orientation and can only grow in the 5’–3’ direction, the mechanism of DNA duplication for two strands are inherently different. Bacteria and viruses use the same processive DNA polymerase for both lagging and leading strands [4]. However, in eukaryotes, a division of labor among the DNA polymerases has been proposed [1, 5]. Extensive genetic studies mostly in yeast suggest that while Polδ plays a key role in lagging strand synthesis, similarly Polε plays an important role in leading strand synthesis [6]. Polδ also takes part in leading strand synthesis in certain contexts and positions. For example, in a yeast strain that lacks Polε catalytic activity, Polδ carries out leading strand synthesis [7]. Also during homologous recombination, re-initiation of replication by DNA Polδ ensures cell survival by replicating both leading and lagging strands simultaneously [8]. Thus, accurate and processive DNA synthesis by Polδ is essential for the preservation of genomic integrity and for the suppression of mutagenesis and carcinogenesis [9]. Therefore, it is important to understand the mechanism underlying processive DNA synthesis by Polδ and decipher precise contribution by its subunits in such an activity.

PCNA, the homotrimeric DNA clamp tethers Polδ to the chromosomal DNA and regulates its function during DNA replication, repair, and recombination [10, 11]. Each PCNA monomer consists of two topologically identical globular domains connected by an interdomain connecting loop (IDCL)[12]. The interaction of PCNA-binding proteins with PCNA is mediated by a conserved PCNA-interacting protein motif (PIP-box) with a consensus sequence Q-x-x-(M/L/I)-x-x-FF-(YY/LY); x being any amino acids [13, 14]. Previously, we have shown that all the three subunits of ScPolδ; Pol3, Pol31, and Pol32 functionally interact with trimeric PCNA mediated by their PIP motifs [15]. To achieve higher processivity *in vitro*, all three PIP boxes are required; however for the cellular function of ScPolδ, along with PIP of ScPol32, one more PIP box from either Pol3 or Pol31 subunit are essential. Interestingly, human Polδ consists of the catalytic large subunit p125 (PolD1) and accessory subunits p50 (PolD2), p68 (PolD3), and p12 (PolD4) [16, 17]. Recently, we have shown that p12 exists as a dimer both in solution as well as in the holoenzyme [16]. In several studies, biochemical interaction between each subunit of hPolδ and PCNA has been demonstrated [17–19]. We showed that the oligomerization of p12 at the amino-terminal domain facilitates its interaction with PCNA at the carboxyl-terminal domain. Both the oligomerization and the PIP motifs of p12 have been mapped [16]. The co-crystal structure of a peptide containing the PIP motif of p68 with PCNA has been solved [20]. Similarly, a PIP motif has been identified in the p50 subunit of hPolδ. Far Western analysis and competition experiment suggested that a 22-mer peptide containing the PIP (_58_LIQMRPFL_65_) of p50 binds PCNA and also competes with full-length p50 for binding to PCNA. Since the binding of p50 to PCNA was inhibited by p21, a yet another PCNA interacting protein, it was suggested that the binding sites for both the proteins are mutually exclusive and p21 has a higher affinity than the p50 [21]. However, there are contradicting reports regarding the interaction of PCNA with the catalytic subunit p125 [21, 22]. One report suggested that p125 alone could not directly interact with PCNA whereas subsequent reports ruled out such a result by carrying out enzymatic assays. Nevertheless, a physical interaction between p125 and PCNA is yet to be established. Therefore, in this study, by using multiple approaches, we report direct physical binding of p125 with PCNA, as well as mapping of their corresponding binding sites. We have also reconfirmed the existence of a multi-subunit human Polδ’s interaction with PCNA, as has been demonstrated previously in *S. cerevisiae* Polδ.

## 2. Results

### 2.1 Sub-cellular co-localization of PCNA with hPolδ subunits

In order to confirm the interaction of hPolδ holoenzyme with PCNA, subcellular localization of individual subunits with PCNA was carried out. In the nucleus, replisomes are microscopically perceived as distinct foci. These foci are considered as chromosomal DNA associated large protein complexes functionally active during DNA replication and other DNA transaction processes. Since PCNA functions as a docking platform, DNA polymerases and other accessory factors accumulate with PCNA in foci [23]. Therefore, whether hPolδ subunits localized to such foci, these subunits were fused to amino-terminal RFP, and PCNA was tagged to GFP. CHO cells were co-transfected with various RFP-fusion constructs viz. p125-RFP, p50-RFP, p68-RFP, p12-RFP and GFP-PCNA. Co-transfectants were fixed and observed under the confocal microscope. As depicted in Fig. 1, while each of the Polδ subunits developed red foci, PCNA formed distinct green punctates in the nucleus. Subsequent merging of both the foci in the co-transfectants led to the appearance of yellow punctuates. Co-localization of foci revealed that p125 (i), p50 (ii), p68 (iii), and p12 (iv) subunits of hPolδ and PCNA functionally interact and develop yellow foci.

**Fig. 1:**
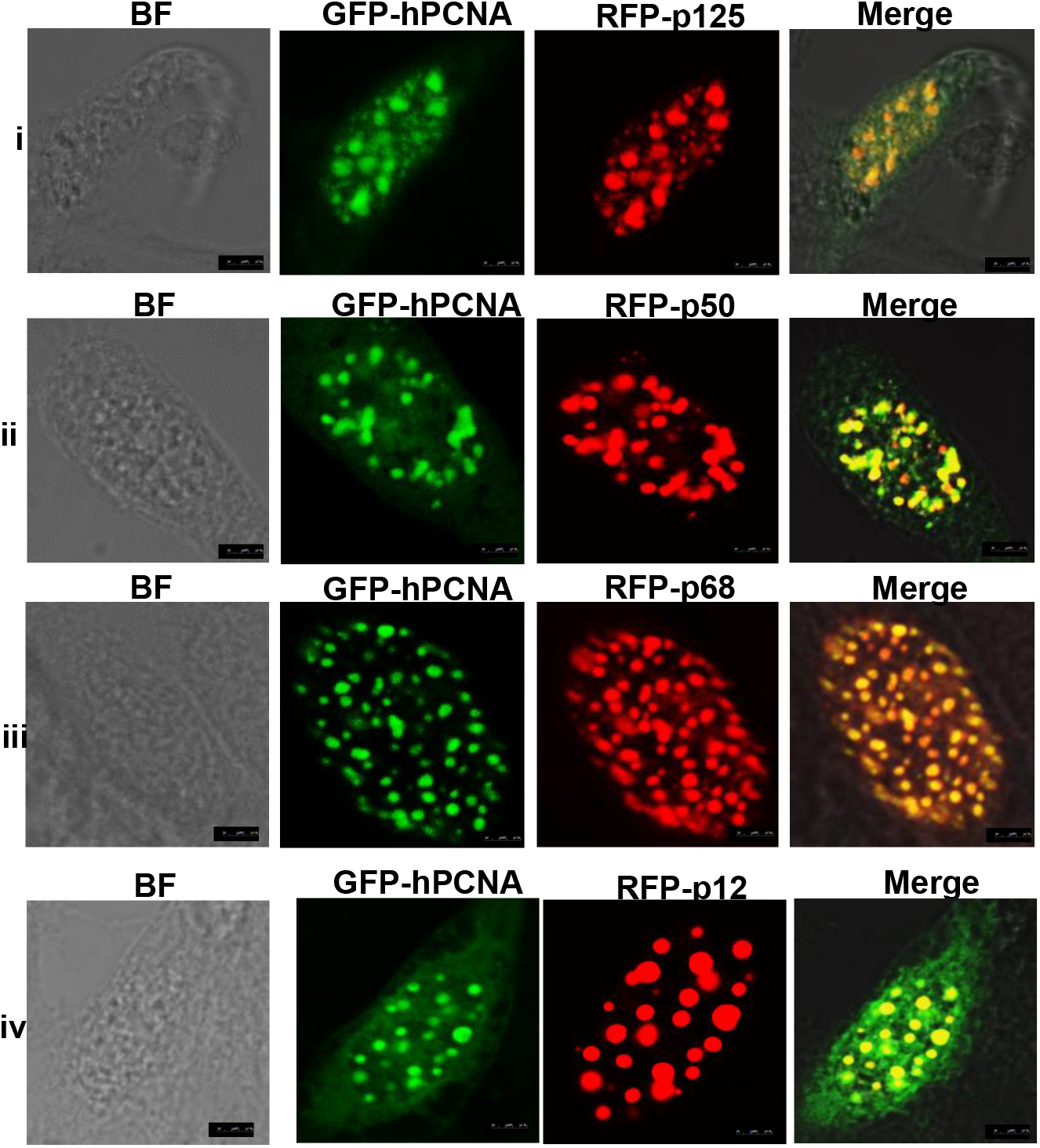
Nuclear co-localization of p125/p50 /p68/p12 with PCNA: CHO cells were transiently co-transfected with GFP-PCNA/RFP-p125 (i), GFP-PCNA/RFP-p50 (ii), GFP-PCNA/RFP-p68 (iii), and GFP-PCNA/RFP-p12 (iv) constructs. After 48 hrs, cells were fixed and mounted and visualized using confocal laser scanning microscopy [Leica TCS SP5] at 63 X objective. Scale bars represent 25 μm.

### 2.2 Identification of putative PIP motifs in p125 subunit of hPolδ

The PIP motif of ScPol3 is located in the vicinity of cysteine-rich metal-binding motifs (CysA and CysB) at the carboxyl-terminal region [15]. At a similar position, we identified a stretch of eight amino acid sequences from 999 - 1006 amino acids in p125 orf (_999_VGGLLAFA_1006_) which shows maximum similarity with the PIP motif in ScPol3 (_996_KGLMSFI_1003_) (Fig. 2 A I). The PIP sequence appears to be highly conserved in other vertebrates Polδ subunits as well (Fig. 2 A II). The crystal structure of the PIP box of p68 (_456_QVSITGFF_463_) and modeled the structure of PIP of p12 (_98_QCSLWHLY_105_) exhibit remarkable structural conservation and form 3_10_ helices [16]. However, according to the p50-p68 co-crystal structure, the PIP motif of p50 (_58_QMRPFL_65_) is located in the α2 helix of the OB-fold domain and is not involved in an interaction with p68 subunit [24]. Although various structures of Polδ from *S. cerevisiae* are available, the putative PIP of Pol3 is not to be visualized, as this portion of the protein is highly unstructured [25, 26]. Therefore, we determined the model structures of putative PIP of Pol3 and p125 and compared them with already confirmed PIPs. The amino acid stretch of PIP sequences from p125 (_996_TGK**V**GG**LL**AFAKR_1008_) and ScPol3 (_993_GSQKGGLMSFIKK_1005_) were considered for peptide structure prediction by using PEP-FOLD3 server (http://bioserv.rpbs.univ-paris-diderot.fr/services/PEP-FOLD3/) without taking any known template to avoid biased prediction (Fig. 2 B). Further, the models were validated Ramachandran plot, which showed most of the residues in allowed regions. Our structural prediction suggested that as in p12 and p68, both the PIPs form a typical 3_10_ helix (I) and superimposes quite nicely with the structure of p68-PCNA co-crystal structure (II). The model structure also predicts the binding of p125-PIP to the IDCL domain of PCNA (Fig. 2B II).

**Fig. 2:**
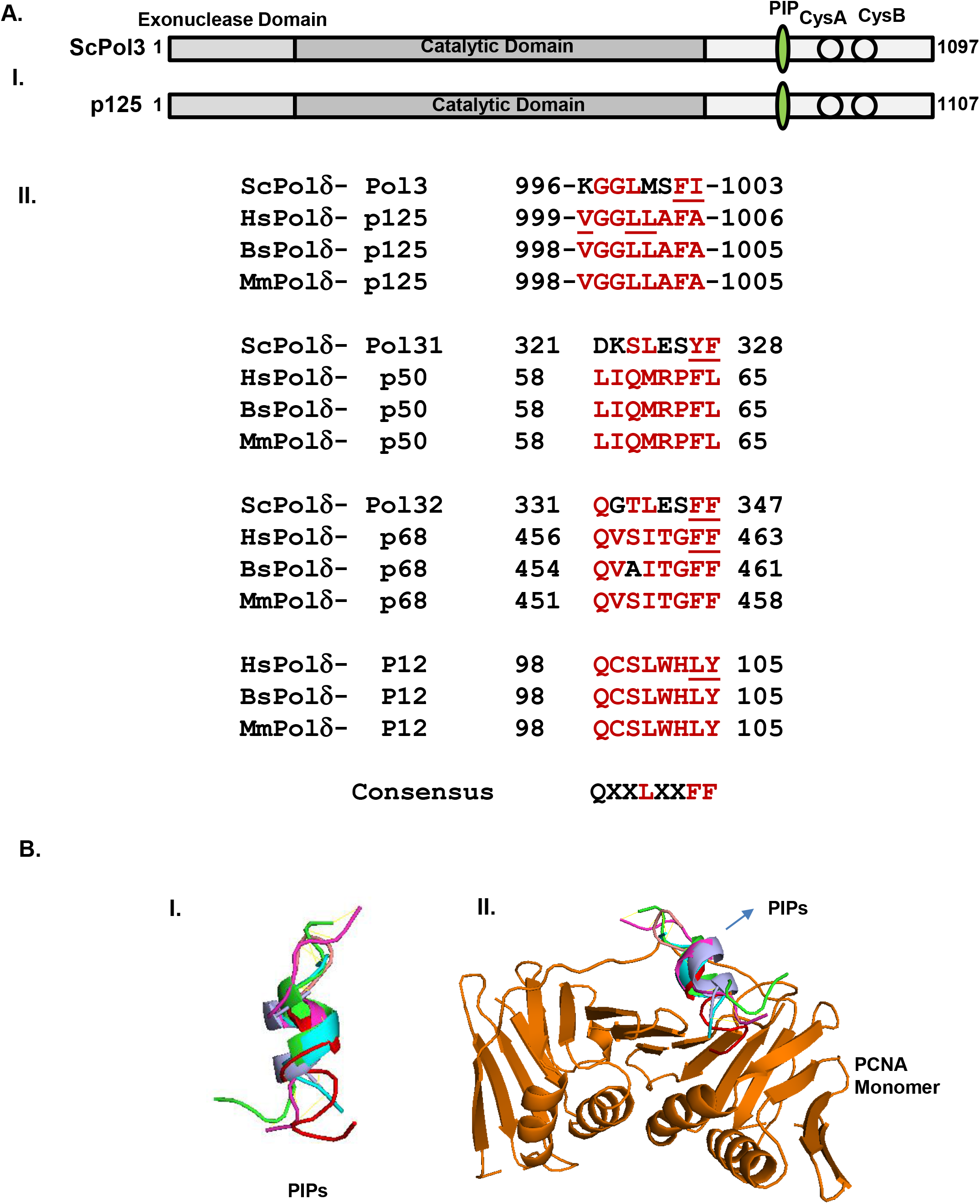
Identification of PIP motifs in Polδ subunits and in silico structure prediction. **A.** Ray diagram representation of various domains of Pol3 and p125 and showing similar laction of PIP motis (I). PCNA interacting motifs in p125, p68, p50 and p12 subunits of Polδ. Amino acid residues in PIP boxes of various Polδ subunits from human (*Homo sapiens*) mouse (*Mus musculus*), bovine (*Bos Taurus*) and budding yeast (*S. cerevisiae*) were aligned. Underlined residues have been mutated (II). **B.** Structure modeling of p125/ScPol3 PIP. Peptide model generated for p125 PIP (_996_TGKVGGLLAFAKR_1008_) and ScPol3 PIP (_993_GSQKGGLMSFIKK_1005_) using PEP-FOLD3 and aligned on already known PIP of p68, p12 and p21 with (i) and without PCNA (ii). PCNA monomer, Pol3, p125, p68, p12 and p21 are shown in orange, cyan, red, green, light blue and purple colors, respectively.

### 2.3 Physical interaction of p125 with hPCNA

To validate the *in silico* prediction of interaction of the putative PIP motif of p125 with PCNA*, in vivo* interaction analyses were carried out by using yeast two-hybrid and co-immunoprecipitation (Fig. 3). Since the aromatic and hydrophobic residues of PIP motifs bind directly to the IDCL of PCNA and mutation in these residues in ScPol3 abrogated its functional interaction with PCNA, we replaced V999, L1002 and L1003 residues of p125 and F462, F463 of p68 with alanines by site-directed mutagenesis. Both wild type and their respective mutants of various subunits of Polδ were fused to the GAL4 binding domain and hPCNA was fused to the GAL4 activation domain. The HFY7C yeast reporter strain was co-transformed with a combination of GAL4 activation and binding domain fusions and transformants were selected on minimal media lacking leucine and tryptophan. The interactions of PCNA with subunits of Polδ in these transformants were analyzed by selecting them on plates lacking histidine (Fig. 3 A). As reported earlier, also in this study, we confirmed the interaction between p12 and p68 subunits with PCNA via their respective identified PIP motifs (compare rows 2 with 3 and 4 with 5). We did not observe any growth when the yeast cells only possessed AD-PCNA and BD empty vector (row 1). In the same assay, we could also observe the growth of the co-transformant of AD-p125 and BD-PCNA on the media lacking histidine suggesting that p125 also interacts with PCNA. However, the respective p125 PIP mutant (V999A/L1002A/L1003A) did not support the growth of the co-transformant with BD-PCNA (compare rows 6 with 7). This result suggests that _999_VGGLLAFA_1006_, _456_QVSITGFF_463_, and _98_QCSLWHLY_105_ are the bonafide PIP motifs of p125, p68 and p12, respectively. PIP of p50 (_58_QMRPFL_65_) has been confirmed by another study. Thus, all the four subunits of human Polδ holoenzyme individually interact with PCNA and their respective PIP motifs were now defined.

**Fig. 3.**
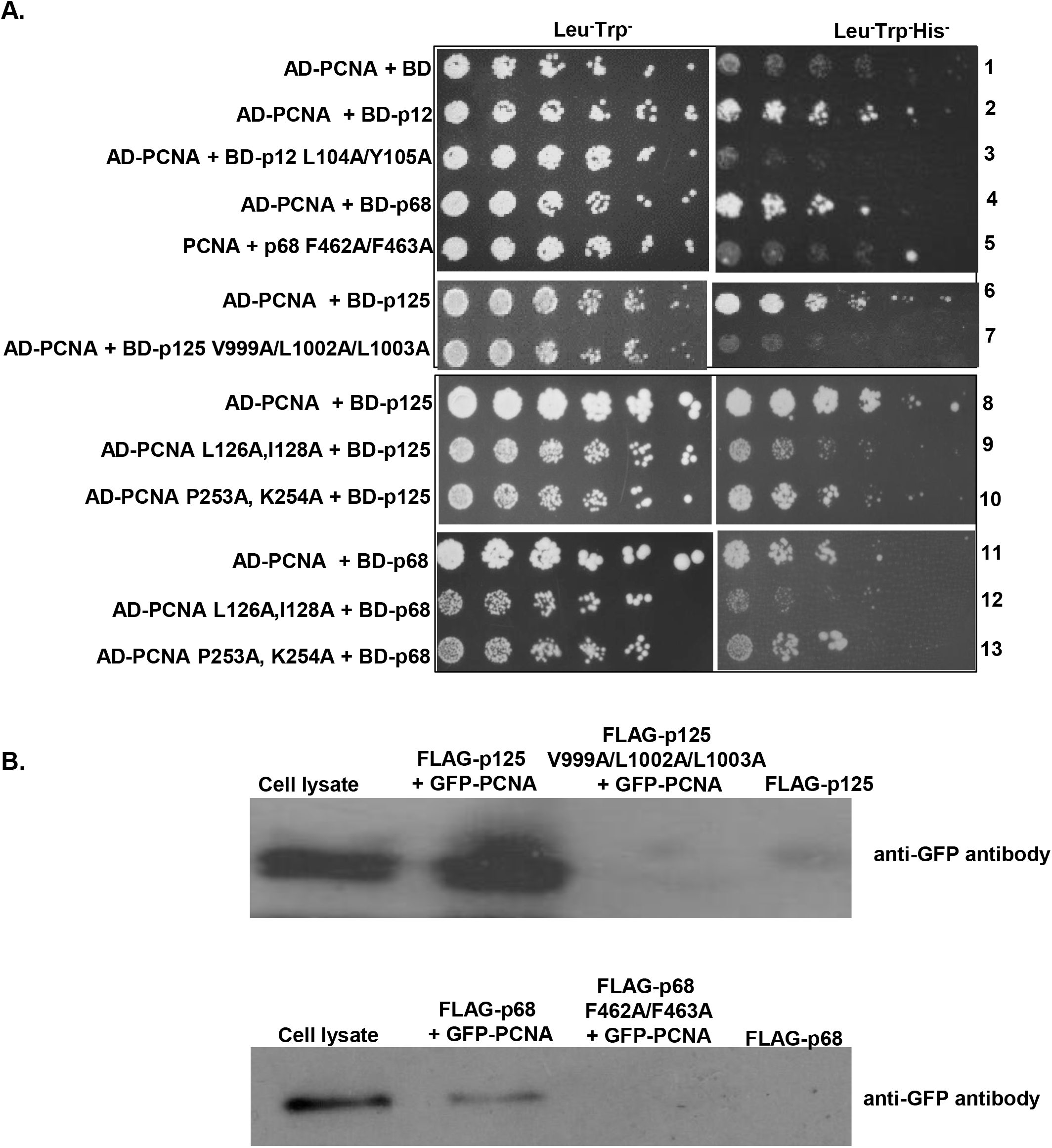
Validation of PIP motifs by yeast two-hybrid and co-immunoprecipitation: **A**. Yeast two-hybrid analysis showing the interaction of PCNA with wild type and putative PIP mutants of p125/p68/p12. HFY7C yeast transformants with various GAL4-AD and BD fusions were selected on SD media plates lacking leucine, tryptophan with and without histidine amino acids. Row 1: AD-PCNA + BD; Row 2: AD-PCNA + BD-p12; Row 3: AD-PCNA + BD-BD-p12 L104A/Y105A; Row 4: AD-PCNA + BD-p68; Row 5: AD-PCNA+ BD-p68 F462A/F463A; Row 6: AD-PCNA + BD-p125, Row 7: AD-PCNA + p125 V999A/L1002A/L1003A; Row 8: AD-PCNA + BD-p125; Row 9: AD-PCNA L126A,I128A + BD-p125; Row 10: AD-PCNA P253A, K254A + BD-p125; Row 11: AD-PCNA + BD-p68; Row 12: AD-PCNA L126A,I128A + BD-p68; and Row 13: AD-PCNA P253A, K254A + BD-p68. **B.** Co-immunoprecipitation study of p125 and p68 with PCNA. HEK cells were transfected with plasmids to express GFP-PCNA, FLAG-p125 or FLAG-p125 PIP mutant, and FLAG-p68 or FLAG-p68 PIP mutant. The co-transfected cells were incubated for 48 hrs and whole-cell lysates (WCL) were prepared. WCL was subjected to Western blotting with an anti-GFP antibody. The PIP mutants in of p125 and p68 were unable to pull down the GFP-PCNA.

To validate further in a homologous expression system, HEK293 cells were cotransfected with GFP-PCNA and wild type or PIP mutants of FLAG-p125 or FLAG-p68 constructs. Using anti-FLAG agarose beads, p125 or p68 proteins were precipitated from the cell lysate. The beads were washed thoroughly and the pull-down of PCNA was detected by probing with anti-GFP antibody (Fig. 3 B). As depicted in the figure, while PCNA was co-precipitated with wild type p125 and p68, their respective PIP mutants failed to pull down PCNA. This result also suggests that both p125 and p68 bind to PCNA in the human cell as well.

### 2.4 p125 binds to inter-domain connecting loop of hPCNA

Inter-domain connecting loop (IDCL) and of the carboxyl-terminal domain of *Saccharomyces cerevisiae* PCNA are the key functional interaction regions of yeast replicative DNA polymerases [27]. Mutational analyses suggested that PCNA with I126A, L128A mutations was defective in interaction and DNA replication by ScPolδ, whereas PCNA with PK252,253AA mutations near the carboxyl terminus was defective in physical and functional interaction with Pol epsilon. Like PIPs of p68 and p12, our predicted model structure also showed p125 PIP binding to the IDCL domain of hPCNA. To confirm the interaction, yeast two-hybrid assay was carried out with p125 or p68 fused to the Gal4 binding domain and two PCNA mutants namely, hpcna-79 and hpcna-90 fused to Gal4 activation domain. hpcna-79 harbors L126A and I128A mutations in IDCL, and hpcna-90 possesses P253A and K254A mutations in the extreme C-terminal tail of PCNA. As depicted in Fig. 3, while the wild-type hPCNA and hpcna-90 were able to interact with both p125 and p68 as evident from the growth on SDA plate lacking leucine, uracil, and histidine (sectors 8, 10, 11 and 13); hpcna-79 did not support the survival as it failed to form intact Gal4 by interacting with p125 or p68 (sector 9 and 12). It suggests that even in humans, Polδ interacts with the IDCL region of PCNA.

### 2.5 Determination of binding affinities of subunits of hPolδ with PCNA by Surface Plasmon Resonance (SPR)

Alignment of PIP boxes in various subunits of Polδ indicated that except PIP in p68 other PIPs are appeared to be non-canonical that lacks the conserved phenylalanine residues (Fig. 2A). The affinity of PIP sequences depends on how accurately the 310 helices fit into the IDCL of PCNA and binding with the neighboring residues [20]. Therefore, we decided to estimate the binding affinities of Polδ subunits with PCNA by using SPR. In this analysis, subunits were passed over the immobilized hPCNA on a GLC-chip (Fig. 4). A concentration-dependent increase in the response unit (~90 RU) was observed when p125 bound to hPCNA. The interaction of p68 and hPCNA was comparatively stronger than the p125 subunit and a good increase in refractive index (~600 RU) and slow dissociation was observed when p68 was bound to PCNA. The *K_D_* values of p68 and p125 complexes with PCNA was determined to be ~7 nM and 100 nM. In similar assay conditions, we did not find any significant binding of p12 and p50 with PCNA by SPR analysis (data not shown). However, in our earlier study, we have determined the affinity equilibrium constant of p12-PCNA by ITC analysis and is found to be in the range of 146 nM [16].

**Figure 4:**
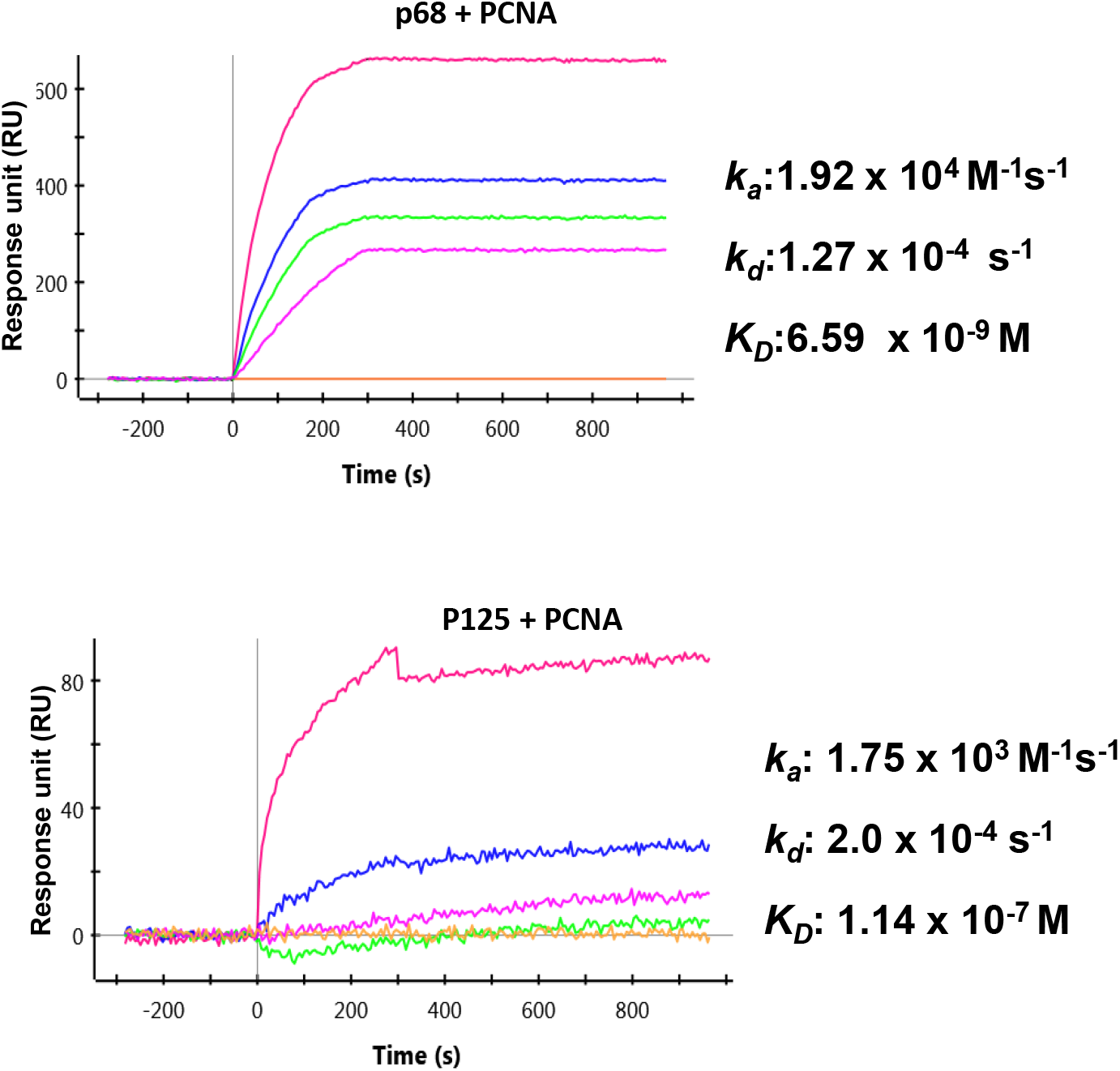
Binding kinetic determination by Surface plasma resonance: Kinetic parameters of interaction between p125 or p68 and PCNA were determined using SPR. Human PCNA was immobilized on GLC chip and various Polδ subunits were passed on it with running buffer at a flow rate of 5Oμl/min for 180s with a 600s dissociation phase. The dissociation constants were determined.

## 3. Discussion

Polδ is the major polymerase involved in DNA synthesis during replication, repair and recombination processes [9]; play a pivotal role in genomic integrity maintenance. Apart from the catalytic subunit, the structural subunits of Polδ also play a crucial role in its fidelity and processivity. Several mutations in the mouse and human Polδ subunits have been mapped to involve in various cancers development [28–30]. The objective of this study was to understand the role of each subunit of hPolδ especially the catalytic subunit in PCNA interaction. Since, hPolδ is a pentameric complex and PCNA is a trimer, the mode of interaction between the two proteins is very complex. Using multiple approaches, we determined the interaction of each subunit with PCNA. By utilizing transiently expressed fluorescently-tagged Polδ subunits and PCNA, we used confocal microscopy to study nuclear co-localization of Polδ subunits with PCNA. Our yeast two-hybrid and Co-IP investigations reveal interactions between PCNA and Polδ subunits viz. p125, p68 and p12. Further, the PIP motifs in each of the subunits were mapped. Similar to ScPol3, the PIP motif of p125 is located in the C-terminal domain upstream to the zinc finger motifs and mutations in this motif abrogated PCNA interaction. Luo X. et al has already mapped the PIP motif in p50 [21]. The co-crystal structure of a peptide possessing PIP motif of p68 with PCNA has already been solved [20] and this study substantiated the earlier reports. In our earlier study, we have shown that p12 is a dimer in hPolδ, and the dimerization in the N-terminal RKR-motif induces its interaction with PCNA via PIP motif located at the C-terminal domain [16]. Our *in silico* structural modeling studies confirm that these motifs form a typical 3_10_ helix and that stably fit into the hydrophobic pocket in the IDCL of PCNA. Taken all together, we suggest that human Polδ interacts with PCNA via multiple identified PIP motifs, and presence of these multiple PIPs in Polδ holoenzyme will stabilize Polδ’s binding to PCNA that in turn will help in processive DNA replication. Considering that improper DNA replication due to altered Polδ interaction with PCNA or its activity can affect genomic stability, our study will help future investigations on Polδ functions.

## 4. Experimental procedures

### 4.1 Generation of various expression constructs for Polδ subunits

Various constructs for the wild type or mutant human PCNA for GST-affinity tag purification or yeast two-hybrid analysis or confocal study have been described previously [16]. Most of the constructs of p125, p68 and p12 used in this study have been described earlier except the site-directed mutants. Site-directed mutagenesis was performed to generate various PIP mutants in p125 and p68. NAP249 (5’-GGT GCT CAC GGG CAA GGC GGG CGG CGC TGC AGC CTT CGC CAA ACG-3’) -NAP250 (5’-CGT TTG GCG AAG GCT GCA GCG CCG CCC GCC TTG CCC GTG AGC ACC-3’) primer pair were used to mutate V999A/L1002A/L1003A in p125 and NAP263 (5’-CCA AAT GAG ACC AGC TGC GGA GAA CCG GGC CCA GC-3’) -NAP264 (5’-GCT GGG CCC GGT TCT CCG CAG CTG GTC TCA TTT GG-3’) primers to mutate F462A/ F463A in p68 by inverse PCR approach. After authenticating their sequence, these orfs were further subcloned into pGAD424.

### 4.2 Yeast two-hybrid analyses

The yeast two-hybrid analyses were performed using HIS3 as a nutritional reporter system as described before [16, 31]. Briefly, the HFY7C yeast strain was transformed with various combinations of the GAL4–AD-PCNA (TRP1) with –BD (LEU2) fusion constructs such as BD-p125, BD-p68, BD-p12, BD-p125 V999A/L1002A/L1003A, BD-p68 F462A/F463A and BD-p12 L104A/Y105A and selected on synthetic dropout media without leucine and tryptophan. To verify interaction, co-transformants were spotted on Leu-Trp-His-selection media plates and incubated further for 2 days at 30 °C before photographed. Yeast transformants exhibiting growth on plates lacking histidine suggest positive protein-protein interaction.

### 4.3 Confocal Microscopy

Chinese hamster ovary (CHO) cells were seeded onto glass coverslips and cultured in standard cell culture conditions as described before [16]. These cells were co-transfected with GFP/RFP fusion constructs of Polδ subunits and using Lipofectamine 3000 transfection kit (Invitrogen). Further, cells were incubated at 37 °C with 5 % CO_2_ and 95 % relative humidity for 2 days. After washings with DPBS, cells were fixed in methanol at −20 °C for 20 mins and again rinsed with DPBS. The coverslips were mounted using antifad and images were taken with Leica TCS SP5 at 63 X objective.

### 4.4 Protein Purifications

All the GST tagged proteins were expressed in either *Escherichia coli* BL21 DE3 under T7 promoter or in YRP654 *S. cerevisiae* under Gal4PGK promoter, and purified by affinity chromatography using glutathione sepharose beads (GE-healthcare). Culture conditions and purification methodology were as described before [16].

### 4.5 Surface Plasmon Resonance

Interaction of PCNA with p125 and p68 was monitored using a Bio-Rad XPR 36 surface Plasmon resonance biosensor instrument. About 5 μg of human PCNA or BSA (~350 RU) was immobilized on GLC chip by amine coupling method as suggested by the manufacturer’s instructions. Purified Polδ subunits was injected at concentration ranging from 125-2000 nM with running buffer composed of 25 mM HEPES pH 7.5, 10 % glycerol, 200 mM Sodium acetate pH 7.8, 8 mM Magnesium acetate, 1 mM DTT, 0.005 % Tween-20 and 0.2 mg/ml BSA, at a flow rate of 50 μl/min for 180 s with a 600 s dissociation phase. Molecular interaction was carried out at 20 °C. Further, the dissociation constants were determined, after fitting the association and dissociation curves to a 1:1 (Langmuir)-binding model.

### 4.6 Co-immunoprecipitation (Co-Ip)

Co-Ip was carried out using HEK293 cells grown up to 70% confluency in a 10 cm dish containing DMEM media supplemented with 10% FBS and 1 X penicillin-streptomycin antibiotics. These cells were co-transfected with GFP-PCNA with either FLAG-p125 or FLAG-p125 V999A/L1002A/L1003A, FLAG-p68 or FLAG-p68 F462A/F463A mutant by using Lipofectamine 3000 transfection kit. Cells were grown in a humidified CO2 incubator at 37°C. After 48 hrs growth, cell were harvested, washed thrice with DPBS and immediately resuspended in RIPA buffer (50 mM Tris-HCl pH 8.0, 0.5 % Sodium deoxycholate, 1000 mM NaCl, 0.1% SDS, 1mM EDTA, 1mM EGTA, 25 mM Sodium Pyrophosphate, 1 mM β-glycerophosphate, 1 mM sodium orthovanadate and protease inhibitor tablet) and kept for 1 hour at 4 °C rocking condition. Followed by centrifugation at 10000 rpm, the supernatant was collected and protein concentration was determined using the Bradford method. About 500 μg of total protein was incubated overnight with anti-FLAG antibody-conjugated agarose beads. The beads were washed thrice with RIPA buffer and bound proteins were eluted by 40 μl of SDS loading buffer and subjected to 12 % SDS PAGE. The proteins from the gel were transferred to PVDF membrane, followed by incubation of the membrane with 5 % skim milk in PBST for 1 hour at room temperature. The blot was washed thrice with PBST and incubated with anti-GFP antibody (1:5000 dilution, cat# ab290 from Abcam) for 2 hours at RT. Subsequently, after through washings, horseradish peroxidase-conjugated goat anti-rabbit IgG (diluted 1:10000 in PBST, cat# A6154; Sigma-Aldrich) was used to develop the blot.

### 4.7 *In silico* analysis of p125 structure

p125 PIP (_996_TGKVGGLLAFAKR_1008_) and ScPol3 PIP (_993_GSQKGGLMSFIKK_1005_) motifs were used for peptide structure prediction by using PEP-FOLD3 server (http://bioserv.rpbs.univ-paris-diderot.fr/services/PEP-FOLD3/). Further, the generated structural models were aligned with PIP peptide sequences of p12, p21 and p68.

## 5. Acknowledgment

We thank Sitendra Prasad Panda for his technical assistance. We also appreciate Bhabani Shankar Sahoo, Amrita Dalei and Shraddheya Kumar Patel for their help in confocal microscopy, SPR, and structural modeling of various proteins, respectively. Our laboratory colleagues are acknowledged for their thoughtful discussion. PK is a DBT-Senior Research Fellow and Dr. Shweta Thakur is a SERB-N-PDF. This work was supported by the intramural core grant from ILS, Bhubaneswar, India

## Abbreviations

PCNA: Proliferating cell nuclear antigen
PIP: PCNA interacting protein
IDCL: Inter domain-connecting loop
SPR: Surface plasmon resonance
ITC: Isothermal calorimetry
GFP: Green fluorescence protein
RFP: Red fluorescence protein

## References

[1] P.M.J. Burgers, T.A. Kunkel, Eukaryotic DNA Replication Fork, Annual review of biochemistry, 86 (2017) 417–438.

[2] B. Stillman, Reconsidering DNA Polymerases at the Replication Fork in Eukaryotes, Molecular cell, 59 (2015) 139–141.

[3] N. Acharya, K. Manohar, D. Peroumal, P. Khandagale, S.K. Patel, S.R. Sahu, P. Kumari, Multifaceted activities of DNA polymerase eta: beyond translesion DNA synthesis, Current genetics, (2018).

[4] M. O’Donnell, L. Langston, B. Stillman, Principles and concepts of DNA replication in bacteria, archaea, and eukarya, Cold Spring Harbor perspectives in biology, 5 (2013).

[5] S.A. Lujan, J.S. Williams, T.A. Kunkel, DNA Polymerases Divide the Labor of Genome Replication, Trends in cell biology, (2016).

[6] T.A. Kunkel, P.M. Burgers, Dividing the workload at a eukaryotic replication fork, Trends in cell biology, 18 (2008) 521–527.

[7] R.E. Johnson, R. Klassen, L. Prakash, S. Prakash, A Major Role of DNA Polymerase delta in Replication of Both the Leading and Lagging DNA Strands, Molecular cell, 59 (2015) 163–175.

[8] I. Miyabe, K. Mizuno, A. Keszthelyi, Y. Daigaku, M. Skouteri, S. Mohebi, T.A. Kunkel, J.M. Murray, A.M. Carr, Polymerase delta replicates both strands after homologous recombination-dependent fork restart, Nature structural & molecular biology, 22 (2015) 932–938.

[9] Y.I. Pavlov, P.V. Shcherbakova, I.B. Rogozin, Roles of DNA polymerases in replication, repair, and recombination in eukaryotes, International review of cytology, 255 (2006) 41–132.

[10] K. Manohar, N. Acharya, Characterization of proliferating cell nuclear antigen (PCNA) from pathogenic yeast Candida albicans and its functional analyses in S. cerevisiae, BMC microbiology, 15 (2015) 257.

[11] G. Maga, U. Hubscher, Proliferating cell nuclear antigen (PCNA): a dancer with many partners, Journal of cell science, 116 (2003) 3051–3060.

[12] T.S. Krishna, X.P. Kong, S. Gary, P.M. Burgers, J. Kuriyan, Crystal structure of the eukaryotic DNA polymerase processivity factor PCNA, Cell, 79 (1994) 1233–1243.

[13] L. Haracska, N. Acharya, I. Unk, R.E. Johnson, J. Hurwitz, L. Prakash, S. Prakash, A single domain in human DNA polymerase iota mediates interaction with PCNA: implications for translesion DNA synthesis, Molecular and cellular biology, 25 (2005) 1183–1190.

[14] J.H. Yoon, N. Acharya, J. Park, D. Basu, S. Prakash, L. Prakash, Identification of two functional PCNA-binding domains in human DNA polymerase kappa, Genes to cells: devoted to molecular & cellular mechanisms, 19 (2014) 594–601.

[15] N. Acharya, R. Klassen, R.E. Johnson, L. Prakash, S. Prakash, PCNA binding domains in all three subunits of yeast DNA polymerase delta modulate its function in DNA replication, Proceedings of the National Academy of Sciences of the United States of America, 108 (2011) 17927–17932.

[16] P. Khandagale, D. Peroumal, K. Manohar, N. Acharya, Human DNA polymerase delta is a pentameric holoenzyme with a dimeric p12 subunit, Life Sci Alliance, 2 (2019).

[17] M. Lee, X. Wang, S. Zhang, Z. Zhang, E.Y.C. Lee, Regulation and Modulation of Human DNA Polymerase delta Activity and Function, Genes (Basel), 8 (2017).

[18] Y. Wang, Q. Zhang, H. Chen, X. Li, W. Mai, K. Chen, S. Zhang, E.Y. Lee, M.Y. Lee, Y. Zhou, P50, the small subunit of DNA polymerase delta, is required for mediation of the interaction of polymerase delta subassemblies with PCNA, PloS one, 6 (2011) e27092.

[19] A.A. Rahmeh, Y. Zhou, B. Xie, H. Li, E.Y. Lee, M.Y. Lee, Phosphorylation of the p68 subunit of Pol delta acts as a molecular switch to regulate its interaction with PCNA, Biochemistry, 51 (2012) 416–424.

[20] J.B. Bruning, Y. Shamoo, Structural and thermodynamic analysis of human PCNA with peptides derived from DNA polymerase-delta p66 subunit and flap endonuclease-1, Structure, 12 (2004) 2209–2219.

[21] X. Lu, C.K. Tan, J.Q. Zhou, M. You, L.M. Carastro, K.M. Downey, A.G. So, Direct interaction of proliferating cell nuclear antigen with the small subunit of DNA polymerase delta, The Journal of biological chemistry, 277 (2002) 24340–24345.

[22] P. Zhang, J.Y. Mo, A. Perez, A. Leon, L. Liu, N. Mazloum, H. Xu, M.Y. Lee, Direct interaction of proliferating cell nuclear antigen with the p125 catalytic subunit of mammalian DNA polymerase delta, The Journal of biological chemistry, 274 (1999) 26647–26653.

[23] J. Essers, A.F. Theil, C. Baldeyron, W.A. van Cappellen, A.B. Houtsmuller, R. Kanaar, W. Vermeulen, Nuclear dynamics of PCNA in DNA replication and repair, Molecular and cellular biology, 25 (2005) 9350–9359.

[24] A.G. Baranovskiy, N.D. Babayeva, V.G. Liston, I.B. Rogozin, E.V. Koonin, Y.I. Pavlov, D.G. Vassylyev, T.H. Tahirov, X-ray structure of the complex of regulatory subunits of human DNA polymerase delta, Cell cycle, 7 (2008) 3026–3036.

[25] R. Jain, M. Hammel, R.E. Johnson, L. Prakash, S. Prakash, A.K. Aggarwal, Structural insights into yeast DNA polymerase delta by small angle X-ray scattering, Journal of molecular biology, 394 (2009) 377–382.

[26] R. Jain, W.J. Rice, R. Malik, R.E. Johnson, L. Prakash, S. Prakash, I. Ubarretxena-Belandia, A.K. Aggarwal, Cryo-EM structure and dynamics of eukaryotic DNA polymerase delta holoenzyme, Nature structural & molecular biology, 26 (2019) 955–962.

[27] J.C. Eissenberg, R. Ayyagari, X.V. Gomes, P.M. Burgers, Mutations in yeast proliferating cell nuclear antigen define distinct sites for interaction with DNA polymerase delta and DNA polymerase epsilon, Molecular and cellular biology, 17 (1997) 6367–6378.

[28] M.K. Swan, R.E. Johnson, L. Prakash, S. Prakash, A.K. Aggarwal, Structural basis of high-fidelity DNA synthesis by yeast DNA polymerase delta, Nature structural & molecular biology, 16 (2009) 979–986.

[29] T. Flohr, J.C. Dai, J. Buttner, O. Popanda, E. Hagmuller, H.W. Thielmann, Detection of mutations in the DNA polymerase delta gene of human sporadic colorectal cancers and colon cancer cell lines, International journal of cancer, 80 (1999) 919–929.

[30] R.E. Goldsby, N.A. Lawrence, L.E. Hays, E.A. Olmsted, X. Chen, M. Singh, B.D. Preston, Defective DNA polymerase-delta proofreading causes cancer susceptibility in mice, Nature medicine, 7 (2001) 638–639.

[31] N. Acharya, L. Haracska, R.E. Johnson, I. Unk, S. Prakash, L. Prakash, Complex formation of yeast Rev1 and Rev7 proteins: a novel role for the polymerase-associated domain, Molecular and cellular biology, 25 (2005) 9734–9740.

